# Mafb-lineage activation of a short polyalanine (+5) PHOX2B mutation produces severe respiratory dysfunction with preserved postnatal weight gain among survivors in a mouse model of congenital central hypoventilation syndrome

**DOI:** 10.64898/2026.09.12.750943

**Authors:** Nelina Ramanantsoa, Charline Riaud, Amelia Madani, Pauline Andres, Thomas Bourgeois, Jean-Luc Mandaba, Laura Cardoit, Jean Forgue, Nathaly Romero, Manon Perrain, Nevine Antoun, Eleonore Sizun, Maud Ringot, Jorge Gallego, Benjamin Dudoignon, Plamen Bokov, Stéphane Dauger, Marie-Pia d’Ortho, Christophe Delclaux, Muriel Thoby-Brisson, Boris Matrot

**Affiliations:** Université Paris Cité, NeuroDiderot, Inserm U1141, Paris, France; Université de Bordeaux, Institut de Neurosciences Cognitives et Intégratives d’Aquitaine, CNRS, Bordeaux, France; Service de Pneumologie, Allergologie, CRCM Pédiatrique, AP-HP, Hôpital Robert Debré, Paris, France; Service d’Explorations Fonctionnelles Pédiatriques, AP-HP, Hôpital Robert Debré, Paris, France; Service de Médecine intensive-Réanimation pédiatriques, AP-HP, Hôpital Robert Debré, Paris, France; Service de Physiologie-Explorations Fonctionnelles, AP-HP, Hôpital Bichat, Paris, France

**Keywords:** congenital central hypoventilation syndrome, PHOX2B, polyalanine expansion, respiratory control, retrotrapezoid nucleus, postnatal growth

## Abstract

**Rationale:** Congenital central hypoventilation syndrome (CCHS) is most commonly caused by polyalanine repeat mutations in PHOX2B. The *in vivo* consequences of the short five-alanine expansion and the contribution of specific hindbrain lineages to the resulting respiratory phenotype remain poorly understood.

**Objectives:** To characterize the neonatal phenotype caused by the *Phox2b^25Ala/+^*mutation and determine the contribution of the MafB lineage associated with the r5–r6 hindbrain territory.

**Methods:** We generated a conditional *Phox2b^25Ala^* allele and studied mice with constitutive or MafB-lineage activation. Neonatal survival, growth, gastric milk content, ventilation, and hypercapnic responses were assessed *in vivo*. Respiratory-network activity and responses to extracellular acidification were recorded in E18.5 isolated brainstem–spinal cord preparations, and retrotrapezoid nucleus (RTN) development was examined histologically.

**Measurements and Main Results:** Constitutive *Phox2b*^25Ala/+^ pups exhibited high neonatal mortality, markedly impaired weight gain, reduced gastric milk scores, hypoventilation, increased apnea time, and blunted ventilatory responses to CO₂. Embryonic preparations showed a slower respiratory rhythm, an impaired response to acidification, and severe RTN dysgenesis. Activation of the mutant allele in the MafB lineage was also associated with severe respiratory dysfunction, RTN dysgenesis, and early neonatal mortality. In contrast, postnatal weight gain was preserved among surviving MafB-lineage mutants, and the reduction in gastric milk scores was significantly attenuated compared with constitutive mutants.

**Conclusions:** A short PHOX2B polyalanine expansion reproduces major respiratory features of CCHS in mice. Activating the mutant allele in the MafB lineage is sufficient to cause severe respiratory dysfunction and neonatal mortality, whereas postnatal weight gain is preserved among survivors.

## Introduction

Congenital central hypoventilation syndrome (CCHS) is a rare, life-threatening neurodevelopmental disorder of autonomic respiratory control, with an estimated incidence of approximately 1 in 148,000 to 1 in 200,000 live births^1^. Its hallmark is impaired automatic ventilatory control, particularly during sleep, together with absent or markedly reduced responses to hypercapnia and hypoxia. Clinical manifestations usually begin at birth or during early infancy, and most affected patients require lifelong ventilatory support^2^. Variable autonomic, gastrointestinal, and neural crest-related manifestations may also occur^1^. More than 90% of patients carry heterozygous polyalanine repeat mutations (PARMs) in PHOX2B, a transcription factor essential for the development of autonomic and respiratory neural circuits^3–5^. These mutations expand the normal 20-alanine tract and produce genotypes ranging from 20/24 to 20/33. Although disease severity generally increases with expansion length, considerable interindividual variability remains^1,6^. The frequent 20/25 genotype is particularly notable for its incomplete penetrance and broad clinical spectrum, ranging from asymptomatic carriage or late-onset disease to neonatal ventilatory failure^7^.

The *in vivo* consequences of shorter PHOX2B polyalanine expansions remain poorly characterized. Current mechanistic understanding relies largely on the *Phox2b^27Ala/+^* (+7) knock-in mouse model, in which severe retrotrapezoid nucleus (RTN) dysgenesis is associated with loss of central CO₂ chemosensitivity. Mutant pups also exhibit impaired respiratory-network activity and abnormalities of upper-airway motor control, but die within hours of birth, limiting the investigation of later postnatal manifestations^8,9^. Restricting the +7 mutation to rhombomeres 3 and 5 severely impaired RTN-dependent chemosensitivity without reproducing the lethality of the constitutive model, indicating that additional PHOX2B-dependent hindbrain circuits contribute to neonatal survival^10^. Complementary studies of a homozygous LBX1 frameshift variant identified in two siblings with PHOX2B-negative CCHS showed that mice carrying the analogous mutation developed severe hypoventilation and died shortly after birth^11^. Conditional mutants expressing this mutation in the MafB lineage encompassing the r5–r6 hindbrain territory exhibited a similar severe respiratory phenotype and neonatal lethality^12^.

The objective of our study was to determine whether the shorter +5 PHOX2B polyalanine expansion reproduces the respiratory phenotype of CCHS *in vivo*, to characterize its additional postnatal effects, and to assess the contribution of the MafB lineage to these phenotypes. We therefore generated a *Phox2b^25Ala/+^* knock-in model and characterized neonatal survival and body-weight gain, ventilation and the hypercapnic response *in vivo*, respiratory-network activity *ex vivo*, and RTN development. We additionally used MafB-Cre-mediated conditional activation of the mutant allele to determine the contribution of this lineage to the respiratory, survival, and body-weight phenotypes caused by the *Phox2b^25Ala^*expansion.

## Methods

### Animals and ethical approval

All procedures were approved by the relevant institutional animal ethics committees and authorized by the French Ministry of Higher Education and Research (APAFIS #45093 and #54728).

### Generation and maintenance of mutant mouse lines

To generate a mutant line expressing a CCHS-causing five-alanine expansion within the 20-residue polyalanine tract of Phox2b, we flanked the endogenous mouse exon 3 with loxP sites and inserted, downstream of the polyA signal, an exon 3 carrying the five-alanine expansion (*Phox2b^25Ala-cki^*). The targeting vector (Fig. 1) was electroporated into C57BL/6N embryonic stem (ES) cells. Correctly targeted ES-cell clones were used to generate chimeric mice. ES cell manipulation and the generation of chimeric mice were done at the Mouse Clinical Institute (PHENOMIN-iCS, Illkirch, France). Male chimeric founders giving germline transmission were crossed with FlpO deleter mice^13^ to remove the neomycin resistance cassette. *Phox2b^25Ala-cki^* line was maintained on a mixed C57BL/6 **×** DBA/2 (B6D2) genetic background. *Phox2b^25Ala-cki/+^*mice were intercrossed to generate *Phox2b^25Ala-cki/25Ala-cki^* homozygous mice. Upon Cre-mediated recombination, the wild-type Phox2b exon 3 and the associated transcriptional STOP/polyadenylation sequences were excised thus allowing expression of the mutated exon 3 carrying the five-alanine expansion. We generated two mutant mouse models carrying the *Phox2b^25Ala^* mutation.

**Figure 1.**
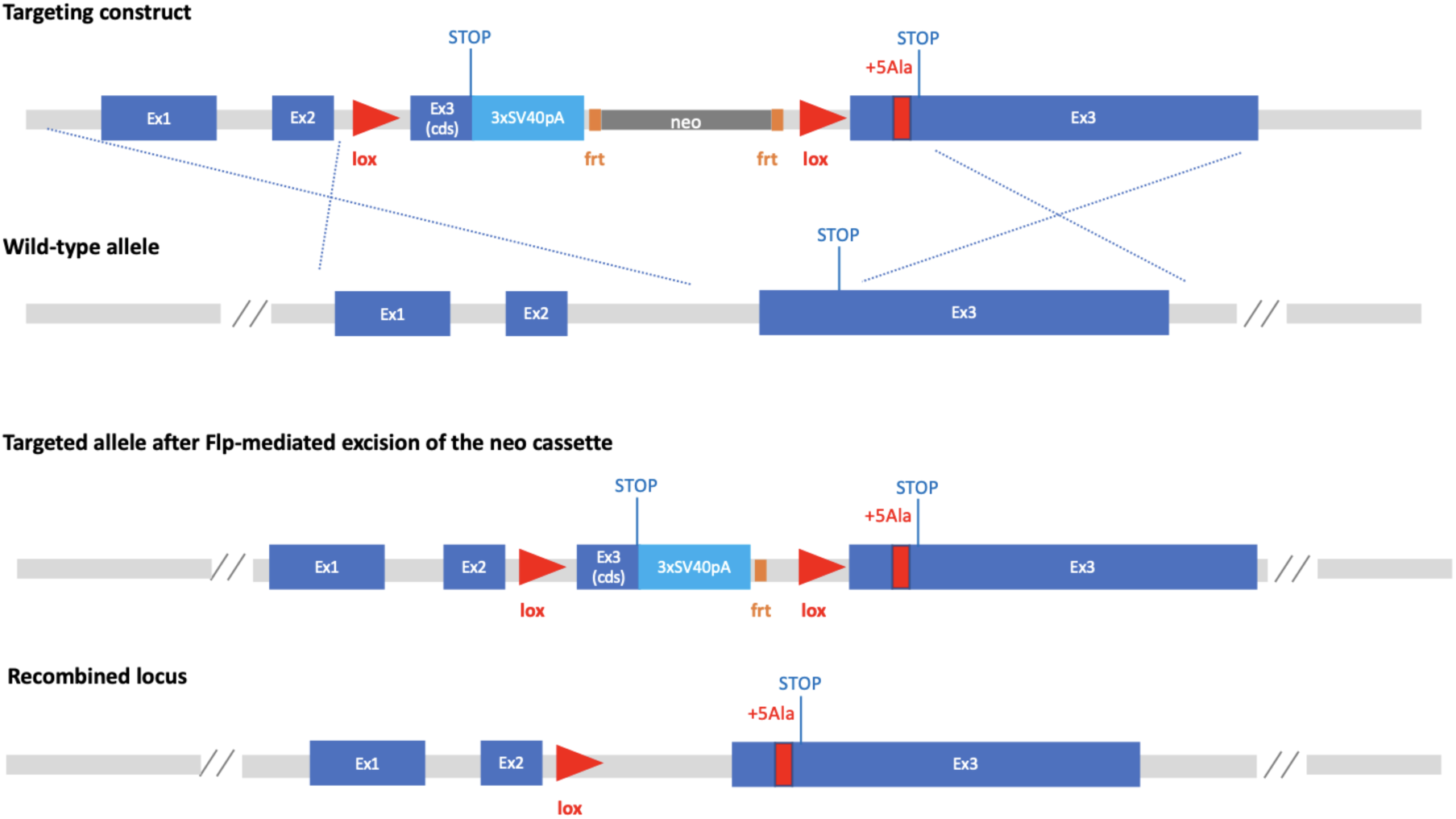
Targeting strategy for the generation of conditional *Phox2b* +5Ala allele. Schematic representation of wild-type *Phox2b* locus and the targeting strategy to conditionally express the +5Ala mutation. The targeting construct contains a wild-type exon 3 coding sequence followed by a 3xSV40 polyadenylation STOP cassette (3xSV40pA), flanked by loxP sites, and downstream an exon 3 carrying the +5Ala expansion. The neo selection cassette, flanked by frt sites, is removed by Flp-mediated recombination. Thus, the resulting targeted allele retains a floxed wild-type exon 3/STOP cassette. Cre-mediated recombination removes the floxed sequence, allowing the expression of mutant +5Ala exon 3 in the recombined locus. Exons in blue, the +5Ala expansion in red, loxP sites as red triangles, frt sites in orange, the 3xSV40pA STOP cassette in cyan and the neo selection cassette in grey. Ex1, exon1; cds, coding sequence; frt, flippase recognition target.

Pups carrying the constitutive mutation were generated by ubiquitous Cre-mediated recombination using the Pgk::Cre deleter line and are hereafter referred to as *Phox2b^25Ala/+^* mice. This strategy was previously used with the conditional *Phox2b^27Ala^* allele to generate constitutive mutants using Pgk::Cre and r3- and r5-lineage mutants using Egr2/Krox20-Cre^10^. Conditional *Phox2b^25Ala^*mutants were generated by crossing *Phox2b^25Ala-cki/+^* mice with *Mafb^Cre/+^* (MafB-mCherry-Cre/+ line)^14^, thereby activating the mutant allele in MafB-lineage cells associated with the r5–r6 hindbrain territory. These mutants are hereafter referred to as *Phox2b^r^*^5*&6-25Ala/+*^ mice. The day on which a vaginal plug was detected was considered as embryonic day 0.5 (E0.5). Genotyping was done on tail DNA using the following primers: (i) to detect the presence of the *Phox2b^25Ala-cki^* allele 5’-GAAGGGCAGACTTGTCGGACTC-3’ and 5’-GCGAAACTTAGCCCGGCG-3’ yielding bands of 223 bp for the wild-type, of 317 bp for the mutated and of 303 bp for the recombined allele, (ii) to detect Cre 5’- AAATTTGCCTGCATTACCG-3’ and 5’-ATGTTTAGCTGGCCCAAATG-3’ yielding a band of 200 bp. Mice were housed in controlled conditions of temperature (21 ± 1°C) and humidity at 55 ± 10% on a 12-h light/dark cycle. Food and water were available ad libitum.

### Experimental design and postnatal phenotyping

Delivery was monitored by continuous video surveillance, and the day of birth was designated postnatal day 0 (P0). *In vivo* phenotyping comprised two complementary protocols applied to constitutive *Phox2b^25Ala/+^* mice, *Phox2b^r5&6-25Ala/+^* mice, and their respective wild-type littermates. Postnatal survival and body weight were monitored from birth to P7, whereas respiratory function was assessed once at P0 by whole-body plethysmography. In both protocols, gastric milk content was estimated at P0 by visually scoring the amount of milk present in the stomach on a three-point scale: 0, no visible milk; 1, intermediate gastric milk content; and 2, milk-filled stomach. Pups of both sexes were studied, and all measurements were performed during the light phase. Except during brief experimental procedures, pups remained with their dam and littermates.

### Postnatal survival, growth, and milk scoring

For longitudinal monitoring, pups were initially assessed 6–12 h after delivery. Each pup was sexed and weighed, and gastric milk content was estimated. Pups were individually identified by marking the back with an indelible marker and were then returned to their home cage. Survival and body weight were subsequently assessed once daily until P7, and individual markings were renewed as required. The time of death was assigned to the first daily assessment at which the pup was found dead. Animals that died during follow-up contributed body weight measurements only up to their last available assessment.

### Whole-body plethysmography

Respiratory measurements were performed 6–18 h after delivery in unanesthetized, unrestrained P0 pups using flow-through whole-body barometric plethysmography, as previously described^10,15^. After sexing, weighing, and scoring of gastric milk content, pups were placed individually in a plethysmography chamber maintained at 32°C. Recordings were performed over 60 min and consisted of 45 min in room air, followed by 5 min of exposure to a hypercapnic gas mixture containing 8% CO₂, 21% O₂, and 71% N₂, and a final 10-min recovery period in room air. Gas was continuously delivered through the chamber at a flow rate of 15 mL/min. Behavior was monitored by video throughout the recording. Activity periods of gross body movement were detected based on video recordings (Logitech C920, Lausanne, Switzerland) and were excluded, and ventilatory variables were quantified during behaviorally inactive periods, which represented approximately 80% of the recording time in all groups. We measured breathing frequency (f_R_, number of breaths/min), tidal volume (V_T_, mL/g), minute ventilation (̇V_E_, calculated as f_R_ × V_T_, mL/min/g), and apneas. Apneas were defined as ventilatory pauses lasting more than 0.90 s, approximately twice the mean respiratory-cycle duration in wild-type pups. Cumulative apnea duration was calculated per minute of recording and is hereafter referred to as apnea time. Baseline ventilatory variables were calculated during behaviorally inactive periods within the first 40 min of the initial 45-min room-air exposure. In every pup, at least 25% of this 40-min window was classified as behaviorally inactive and included in the analysis. Responses to hypercapnia were assessed by comparing values obtained during the last 5 min of this room-air period (immediately preceding CO₂ exposure) with those obtained during the final 3 min of the 8% CO₂ challenge, retained only if at least 10% of the relevant window was classified as behaviorally inactive, since CO₂ exposure elicits a behavioral response that reduces the proportion of usable signal.

### *Ex vivo* brainstem–spinal cord electrophysiology

Electrophysiological recordings of rhythmically organized breathing-related activities were performed on isolated brainstem preparations, which retain the different central respiratory networks. Because mutant newborns die rapidly after birth, recordings were performed at embryonic day (E)18.5. Both recordings and analyses were performed blind to genotype, the genotype of each embryo being determined *a posteriori*. To ensure precise embryonic staging, mice were mated for a single night.

Pregnant mice were killed by cervical dislocation. E18.5 embryos were rapidly removed from the uterine horns by cesarean section and placed in oxygenated artificial cerebrospinal fluid (aCSF) at 18**–**20°C until dissection. Then, embryos were decerebrated and decapitated. The brainstem was carefully separated from surrounding tissues, and completely isolated through a rostral section made at the meso-rhombencephalic limit, and a caudal section made below the fourth cervical roots. Attention was paid to ensure that the ventral cervical motor roots remained intact.

Brainstem preparations were dissected in cold (4°C) artificial cerebrospinal fluid (aCSF) solution composed of (in mM): 120 NaCl, 8 KCl, 0.58 NaH2PO4, 1.15 MgCl2, 1.26 CaCl2, 21 NaHCO3, 30 glucose (pH 7.4) and equilibrated with 95% O_2_**–**5% CO_2_. To mimic hypercapnia *in vitro*, aCSF acidosis (pH 7.2) was induced by decreasing NaHCO3 concentration to 10 mM and increasing NaCl concentration to 130 mM. Isolated brainstem preparations were positioned ventral side-up in the recording chamber and continuously superfused with oxygenated aCSF at 30°C. A recovery period of 30 min was observed before starting any recordings.

All recordings were performed using borosilicate glass electrodes (Harvard Apparatus GC150F, Germany), filled with aCSF solution (see above), broken at the tip to match the diameter of the targeted phrenic root, and connected through a silver wire to a high-gain amplifier (A-M Systems, USA). Signals were filtered (Differential AC amplifier model 1700, bandwidth 100 Hz**–**1 KHz), rectified and integrated (NeuroLog system Digitimer Ltd., England), digitized (Axon Digidata 1440A, Molecular Devices, USA), then recorded and analyzed offline using pCLAMP 10 software (Axon Instruments). Measurements of the respiratory frequency were performed over a 2-minute period in control conditions (pH 7.4) and in response to acidosis (pH 7.2).

### Immunohistochemistry

Isolated brainstem preparations were placed in 4% paraformaldehyde for 1 night to allow tissue fixation. Coronal and sagittal frozen sections were obtained by first placing brainstems overnight in a 20% sucrose-PBS (phosphate-buffered saline) solution for cryoprotection, and then embedding them in a block of Tissue Tek (Leica, Nanterre, France). Thirty-micrometer-thick sections were obtained by cryostat sectioning (CM 3050 S, Leica). To limit nonspecific labeling, preparations were pre-incubated in a solution of PBS containing 0.3% Triton X-100 and 1% BSA for 90 min. Primary antibodies were mouse anti-Phox2b (1/100; ref SC-376997, Santa Cruz), goat anti-Phox2b (1/500; ref AF4940, Biotechne), mouse anti-Islet1,2 (1/250; ref 40.2D6/39.4D5, DSHB), goat anti-ChAT (1/100; ref AB144P, Millipore) and rabbit anti-NK1R (1/10000; S8305, Merck). They were applied overnight at room temperature and under slight agitation. After washing, sections were incubated for 90 min at room temperature with the appropriate secondary antibodies (all 1 :500 ; Thermo Fisher Scientific) : Alexa Fluor 488-conjugated anti-mouse, Alexa Fluor 568-conjugated anti-goat and Alexa Fluor 647-conjugated anti-rabbit antibodies. Stained slices were cover-slipped and mounted in Vectashield Hard Set medium (Eurobio, Les Ulis, France) and kept in the dark until imaging using a slide scanner (VS200, Olympus). Cell counting and measurements were performed on every second section and blindly with respect to genotype. Neurons were counted either manually when possible or semi-automatically using the QuPath software. Genotypes were disclosed only after image acquisition and cell counting had been completed.

### Statistics

Survival distributions were estimated using the Kaplan–Meier method and compared between each mutant group and its wild-type littermates using two-sided log-rank (Mantel– Cox) tests. Animals alive at P7 were right-censored.

Missing body weights located between observed measurements were estimated by linear interpolation; no extrapolation was performed beyond the last available measurement. Longitudinal body-weight data at P0, P2, P4, and P7 were analyzed using linear mixed-effects models, with genotype, age, and their interaction as fixed effects, litter as a random intercept, and animal-specific random intercepts and slopes to account for repeated measurements. Animal-specific random intercepts and slopes were fitted as uncorrelated. Degrees of freedom were approximated using Satterthwaite’s method. Prespecified genotype contrasts were estimated at each age, with *P* values adjusted for the four comparisons using Holm’s method. A complementary model treating age as a continuous variable was used to estimate genotype-specific rates of weight gain and their difference, with 95% confidence intervals. Sensitivity analyses were conducted using observed body weights only.

Within each experimental cohort, milk scores were compared between genotypes using a two-sided exact stratified Wilcoxon–Mann–Whitney test, with litter as the stratification factor. Only litters containing both genotypes contributed to the test.

To compare genotype-associated changes in milk-score distributions between the constitutive and MafB-lineage models, data from the four experimental cohorts were analyzed jointly using a cumulative-link mixed model with a logit link. Milk score was treated as an ordered categorical outcome. Experimental cohort, mutant status, and the mutant-status × model interaction were included as fixed effects, and litter was included as a random intercept. The interaction tested whether the mutant-versus-WT difference differed between the MafB-lineage and constitutive models. Model coefficients are reported as log-odds estimates together with odds ratios and 95% confidence intervals. As a sensitivity analysis, milk score was also treated as a numerical outcome and analyzed using a linear mixed-effects model with the same fixed- and random-effects structure.

Baseline ventilatory variables were compared between genotypes using two-sided unpaired Student’s *t* tests. Ventilatory responses to 8% CO₂ were analyzed using two-way mixed-design ANOVA, with genotype as the between-subject factor and gas condition (room air versus 8% CO₂) as the within-subject repeated-measures factor. The genotype × gas-condition interaction was used to determine whether the response to hypercapnia differed between genotypes. When this interaction was significant, prespecified within-genotype comparisons between room air and 8% CO₂ were performed with Holm-Šídák adjustment for the two comparisons.

Changes in respiratory burst frequency induced by extracellular acidification were analyzed using two-way mixed-design ANOVA, with genotype as the between-preparation factor and pH condition (pH 7.4 versus pH 7.2) as the within-preparation repeated-measures factor. When the genotype × pH-condition interaction was significant, prespecified simple-effects comparisons assessed the effect of acidification within each genotype and the effect of genotype at each pH condition, with Holm–Šídák adjustment for multiple comparisons. RTN neuronal counts were compared between genotypes using a two-sided exact Mann–Whitney test for constitutive *Phox2b^25Ala/+^* embryos and a two-sided unpaired Welch’s *t* test for conditional *Phox2b^r5&6-25Ala/+^*embryos, with the embryo as the statistical unit.

Data acquisition and analysis were performed blinded to genotype whenever feasible. Data from both sexes were pooled because the study was not designed or powered to estimate sex-specific effects. Analyses were performed using Prism 11 (GraphPad Software) and R with the coin, lme4, lmerTest, emmeans, and ordinal packages. All tests were two-sided, and *P* < 0.05 was considered statistically significant. Continuous data are reported as mean ± SEM in the text; graphical summaries and categorical distributions are described in the corresponding figure legends.

Generative AI tools (ChatGPT and Codex, OpenAI) were used to assist with language editing and statistical-code development. All analyses, outputs and manuscript content were reviewed and approved by the authors, who take full responsibility for the accuracy and integrity of the work.

## Results

### Constitutive *Phox2b^25Ala/+^* mice exhibit neonatal lethality and impaired postnatal growth

Postnatal survival and growth were assessed during the first week of life in constitutive *Phox2b^25Ala/+^* mice (n = 18) and wild-type littermates (n = 22) from 5 litters. Survival was markedly reduced in *Phox2b^25Ala/+^* mice compared with wild-type littermates (*P* < 0.001; Fig. 2A), with most deaths occurring within the first 72 hours after birth. Postnatal growth among surviving pups was also markedly impaired (Fig. 2B). Longitudinal analysis revealed a significant genotype × age interaction (*P* < 0.001; Fig. 2C), indicating that body-weight trajectories differed between genotypes. Birth weight was comparable between wild-type and *Phox2b^25Ala/+^*pups (1.38 ± 0.02 g vs. 1.30 ± 0.04 g, respectively; Holm-adjusted *P* = 0.24), whereas surviving mutants had significantly lower body weights at P2 (adjusted *P* = 0.001), P4, and P7 (adjusted *P* < 0.001 for both). The estimated mean rate of weight gain was 0.403 g/day in wild-type pups and 0.083 g/day in surviving mutants, corresponding to a mutant-versus-WT difference of −0.320 g/day (95% CI, −0.432 to −0.208; *P* < 0.001). Analyses restricted to observed body weights yielded similar results. Gastric milk scores at P0 were also significantly lower in *Phox2b^25Ala/+^* mutants than in wild-type littermates after accounting for litter (*P* = 0.001; Fig. 2D). Thus, constitutive *Phox2b^25Ala/+^* mutants had comparable birth weights but showed lower gastric milk content, high neonatal mortality, and markedly impaired postnatal weight gain among survivors.

**Figure 2.**
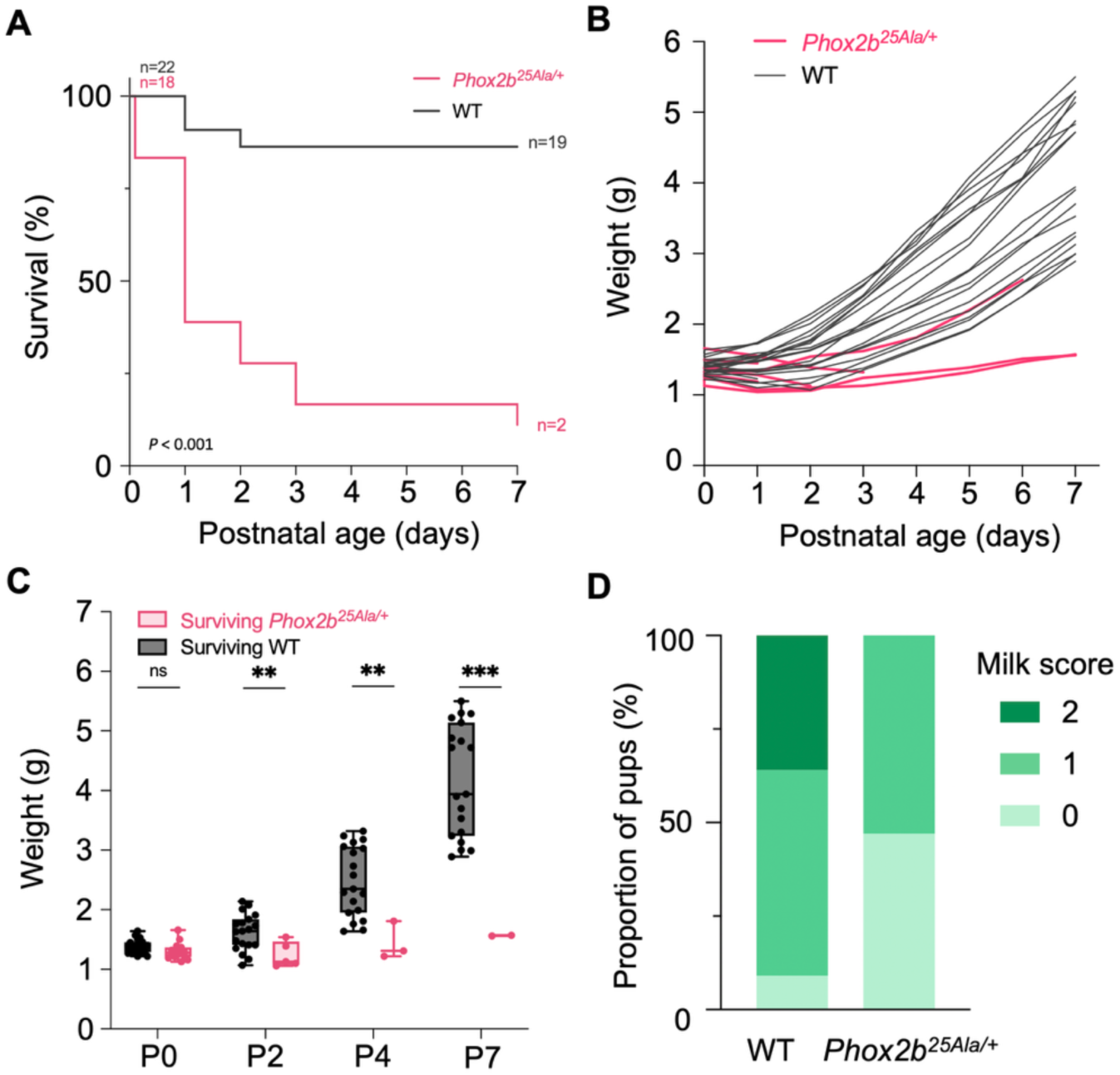
Early postnatal survival and growth of constitutive *Phox2b^25Ala/+^* mice. (A) Kaplan– Meier survival curves of constitutive *Phox2b^25Ala/+^* mice (n = 18, pink) and wild-type (WT) littermates (n = 22, black) from birth (P0) to postnatal day 7 (P7). Three mutant pups were found dead at birth. Survival was markedly reduced in *Phox2b^25Ala/+^* (two-sided log-rank test, *P* < 0.001). (B) Individual body weight trajectories of *Phox2b^25Ala/+^* mice (n = 15, pink) and WT littermates (n = 22, black) from P0 to P7. Individual trajectories are discontinued after death. (C) Body weights of *Phox2b^25Ala/+^* mice and WT littermates alive at each indicated age. Body weight trajectories differed significantly between genotypes (genotype × age interaction, *P* < 0.001), with marked postnatal growth impairment in *Phox2b^25Ala/+^* mice: body weights were comparable between *Phox2b^25Ala/+^* and WT mice at P0 (n = 15 and n = 22), whereas mutant mice had significantly lower body weights at P2 (n = 5 and n = 19), P4 (n = 3 and n = 19), and P7 (n = 2 and n = 19). For all comparisons, sample sizes are reported for *Phox2b^25Ala/+^* and WT mice, respectively. Boxplots show the median, interquartile range, and full range, with individual animals represented by symbols. (D) Distribution of milk scores (0, no visible milk; 1, intermediate gastric milk content; and 2, milk-filled stomach) in wild-type (WT, n =22) and *Phox2b^25Ala/+^* (n =15) pups at P0. Milk scores were lower in *Phox2b^25Ala/+^* pups compared with WT littermates (two-sided exact stratified Wilcoxon–Mann–Whitney test, stratified by litter, *P* = 0.001). Bars show the proportion of pups assigned to each score within each genotype. ** Holm-adjusted *P* < 0.01; *** Holm-adjusted *P* < 0.001; ns, not significant.

### Constitutive *Phox2b^25Ala/+^* mice exhibit neonatal hypoventilation and a blunted ventilatory response to CO₂

Baseline ventilation and ventilatory response to hypercapnia were birth in constitutive *Phox2b^25Ala/+^* mice using whole-body plethysmography (six litters; Fig. 3). In normoxia, *Phox2b^25Ala/+^* mutant pups (n = 10) exhibited marked hypoventilation compared with wild-type littermates (n = 19), with minute ventilation (̇V_E_) reduced by 52% (*P* < 0.01; Fig. 3B). This reduction was primarily driven by a 52% lower respiratory rate (f_R_, *P* < 0.001), whereas tidal volume (V_T_) was similar between groups (*P* = 0.90). *Phox2b^25Ala/+^* pups exhibited a 2.9-fold higher apnea time than wild-type littermates (*P* < 0.001).

**Figure 3.**
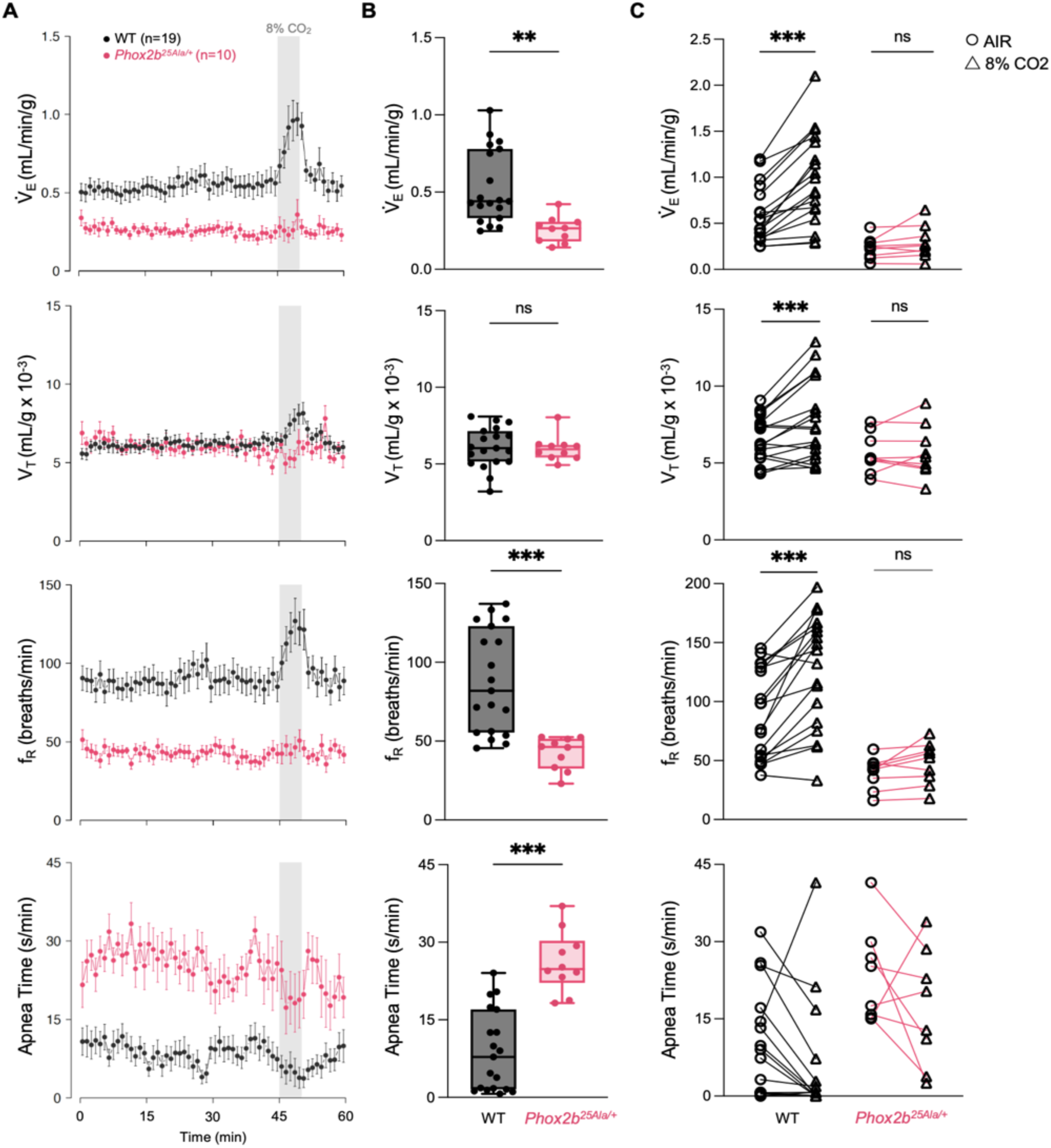
Markedly depressed basal ventilation and blunted ventilatory response to hypercapnia in *Phox2b^25Ala/+^* mutants compared with wild-type (WT) littermates at birth, measured by whole-body plethysmography at 32°C. For each panel, minute ventilation (Vd_E_), tidal volume (V_T_), breathing frequency (f_R_), and apnea time are shown. (A) Ventilatory tracings (each dot represents mean ± SEM over a 1-min period) of *Phox2b^25Ala/+^* mutant pups (n = 10, pink) and their WT littermates (n = 19, black) over the 60-min recording protocol, consisting of 45 min in room air, followed by 5 min of exposure to a gas mixture containing 8% CO₂ (gray shaded area), and 10 min of recovery in room air. (B) Baseline ventilation in *Phox2b^25Ala/+^* mutants (n = 10, pink) compared with WT littermates (n = 19, black). Values were calculated during the first 40 min of exposure to room air. Boxplots show the median, interquartile range, and full range, with individual animals represented by symbols. (C) Individual ventilatory responses to hypercapnia in *Phox2b^25Ala/+^* mutants (n = 9) and WT littermates (n = 18). Values were calculated during the 5 min of breathing room air preceding hypercapnic exposure (circles) and during the last 3 min of hypercapnic exposure (triangles). Paired measurements from individual pups are connected by lines (*Phox2b^25Ala/+^*: pink; WT: black). Two-way mixed ANOVA revealed significant genotype x gas-condition interactions for ̇V_E_ (*P* = 0.002), V_T_ (*P* = 0.04), f_R_ (*P* = 0.007) but not for apnea time (*P* = 0.67). For variables showing a significant interaction, Holm-Šídák-adjusted within-genotype comparisons showed significant increases during exposure to 8% CO₂ in WT pups, whereas the corresponding changes were not significant in *Phox2b^25Ala/+^* pups. No post hoc comparisons were performed for apnea time. Adjusted comparisons are indicated in the graphs. ** *P* < 0.01; *** *P* < 0.001; ns, not significant.

Ventilatory responses to 8% CO₂ differed significantly between genotypes for ̇V_E_ (genotype × gas-condition interaction, *P* = 0.002), f_R_ (*P* = 0.007) and V_T_ (*P* = 0.04) but not for apnea time (*P* = 0.67; Fig. 3C). In wild-type pups (n = 18), CO₂ exposure increased f_R_ by 41%, V_T_ by 19% and ̇V_E_ by 67% (Holm-Šídák-adjusted *P* < 0.001 for all comparisons). By comparison, the corresponding increases of 19%, 2% and 25% in *Phox2b^25Ala/+^*pups were not statistically significant (adjusted *P* = 0.35, *P* = 0.83 and *P* = 0.46, respectively). Apnea time decreased during CO₂ exposure in both genotypes, but the non-significant genotype × gas-condition interaction provided no evidence that the magnitude of this response differed between genotypes. Gastric milk scores observed before plethysmography were lower in *Phox2b^25Ala/+^* pups than in their wild-type littermates (*P* < 0.001; Fig. 4), confirming the previous observation in the survival and growth protocol. Together, these findings show that the +5 PHOX2B expansion produces neonatal hypoventilation, increased apnea time, and a markedly blunted ventilatory response to CO₂, thereby reproducing the cardinal respiratory features of CCHS.

**Figure 4.**
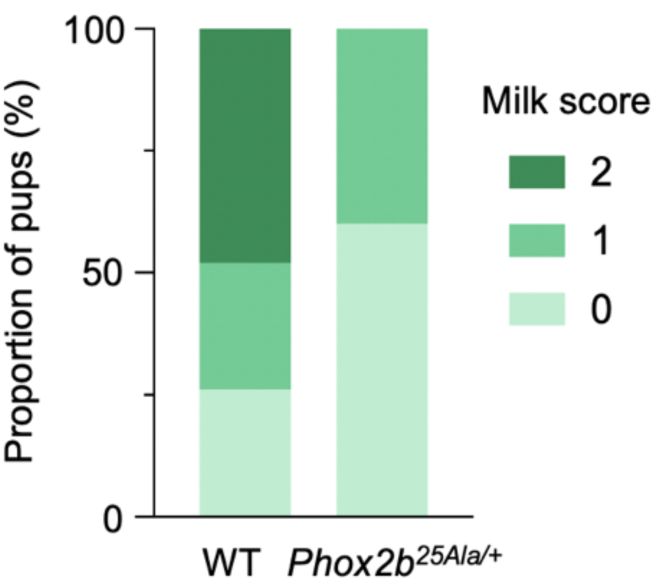
Gastric milk scores before plethysmography recording in wild-type (WT, n = 19) and constitutive *Phox2b^25Ala/+^* mutant pups (n = 10) at P0. Bars show the proportion of pups assigned to each milk score (0, no visible milk; 1, intermediate gastric milk content; and 2, milk-filled stomach) within each genotype. Milk-score distributions were shifted towards lower values in mutants compared with WT littermates (two-sided exact stratified Wilcoxon–Mann–Whitney test, stratified by litter, *P* < 0.001). All pups are shown; the statistical comparison included 12 WT and 10 mutant pups from litters containing both genotypes.

### Constitutive *Phox2b^25Ala/+^* embryos exhibit reduced respiratory rhythm and impaired acidification responses *ex vivo*

Isolated brainstem–spinal cord preparations obtained at embryonic day (E) 18.5 exhibited stable rhythmic phrenic motor activity in both wild-type (n = 12) and *Phox2b^25Ala/+^* (n = 11) embryos (Fig. 5A). However, respiratory burst frequency was lower in mutant than in wild-type preparations (8.20 ± 0.71 vs 11.0 ± 0.93 bursts/min; Holm–Šídák-adjusted *P* = 0.04; Fig. 5B). We next examined the response of the respiratory network to extracellular acidification from pH 7.4 to pH 7.2, a well-established surrogate for central chemosensory stimulation. The response differed significantly between genotypes, as indicated by a significant genotype × pH interaction (*P* = 0.009). Acidification increased respiratory burst frequency in wild-type preparations (adjusted *P* < 0.001), whereas the smaller increase in *Phox2b^25Ala/+^* preparations was not statistically significant (adjusted *P* = 0.08). Consequently, burst frequency was markedly lower in mutant than in WT preparations at pH 7.2 (adjusted *P* = 0.001; Fig. 5B). These findings indicate that the +5 PHOX2B expansion is associated with a slower embryonic respiratory rhythm and a markedly impaired respiratory-network response to extracellular acidification *ex vivo*.

**Figure 5.**
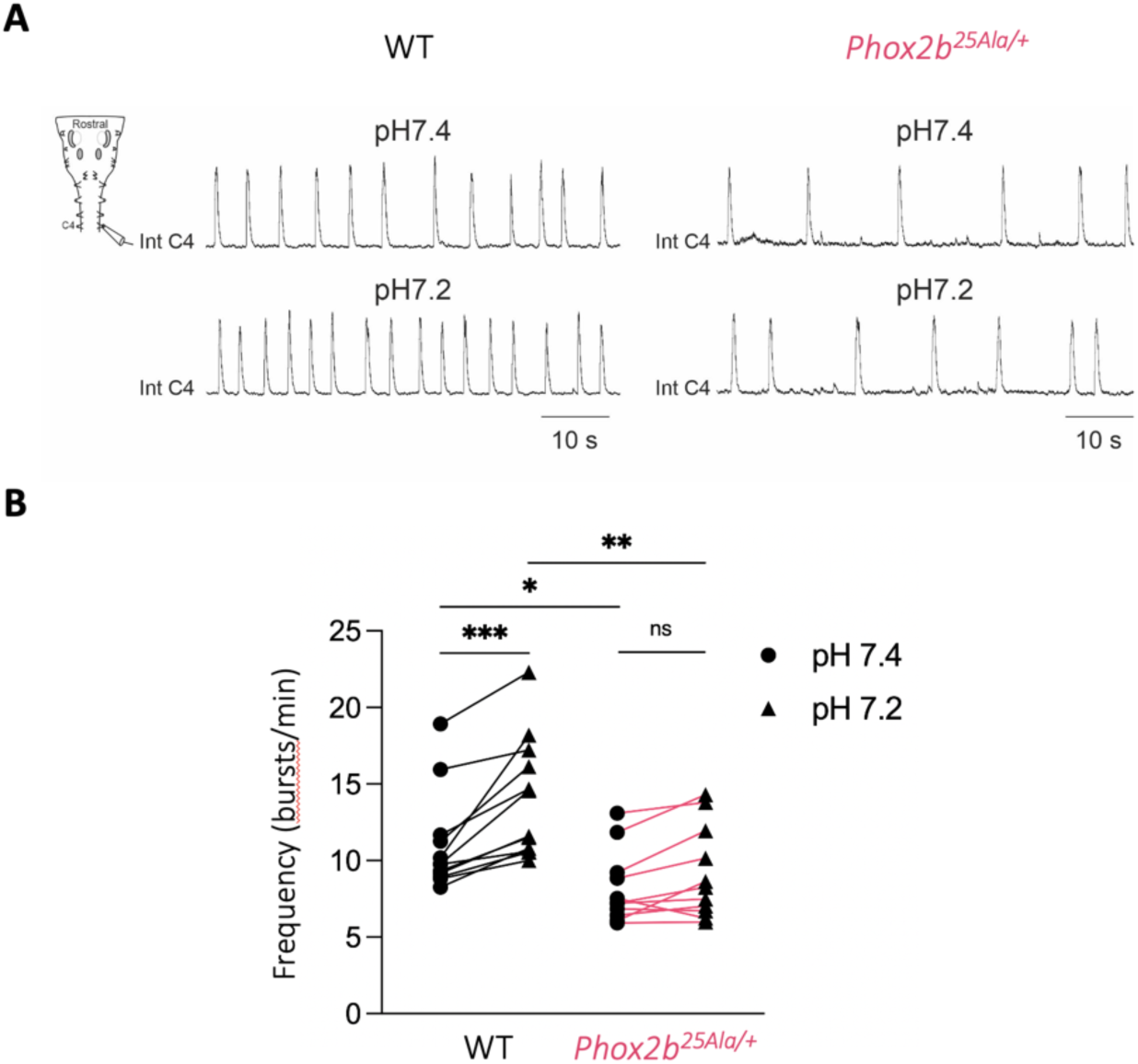
Reduced basal respiratory rhythm and impaired acidification-induced stimulation in isolated brainstem–spinal cord preparations from *Phox2b^25Ala/+^* embryos. (A) Representative integrated recordings of phrenic nerve (C4) activity obtained from E18.5 isolated brainstem– spinal cord preparations from wild-type (WT) and constitutive *Phox2b^25Ala/+^* embryos superfused with artificial cerebrospinal fluid at pH 7.4 and after extracellular acidification (pH 7.2). (B) Mean respiratory burst frequency calculated over 2-min recording periods at pH 7.4 and pH 7.2 in WT (n = 12) and *Phox2b^25Ala/+^* (n = 11) preparations. Individual preparations are connected by lines (WT: black; *Phox2b^25Ala/+^*: pink). Two-way mixed-design ANOVA, with pH condition as the repeated-measures factor, revealed a significant genotype × pH interaction (*P* = 0.009). Holm–Šídák-adjusted comparisons showed that extracellular acidification increased burst frequency in WT preparations (*P* < 0.001), whereas the smaller increase in *Phox2b^25Ala/+^* preparations was not statistically significant (*P* = 0.08). Burst frequency was lower in *Phox2b^25Ala/+^* than in WT preparations both at pH 7.4 (*P* = 0.04) and at pH 7.2 (*P* = 0.001). * *P* < 0.05; ** *P* < 0.01; *** *P* < 0.001; ns, not significant.

### Constitutive *Phox2b^25Ala/+^* embryos exhibit severe RTN dysgenesis

At E18.5, the RTN was markedly affected in constitutive *Phox2b^25Ala/+^*embryos, as previously reported in mouse models carrying the +7 PHOX2B expansion (Fig. 6A,B). RTN neurons, identified as ChAT−/ISLET1,2−/PHOX2B+/NK1R+ cells located ventral to the ChAT+/ISLET1,2+/PHOX2B+ facial motor nucleus, were markedly reduced in mutants compared with wild-type littermates (73 ± 22 versus 538 ± 79 neurons, respectively; two-sided exact Mann–Whitney test, *P* < 0.001; Fig. 6C). Thus, despite its shorter length, the +5 polyalanine expansion is sufficient to severely impair the development of this central chemoreceptor population.

**Figure 6.**
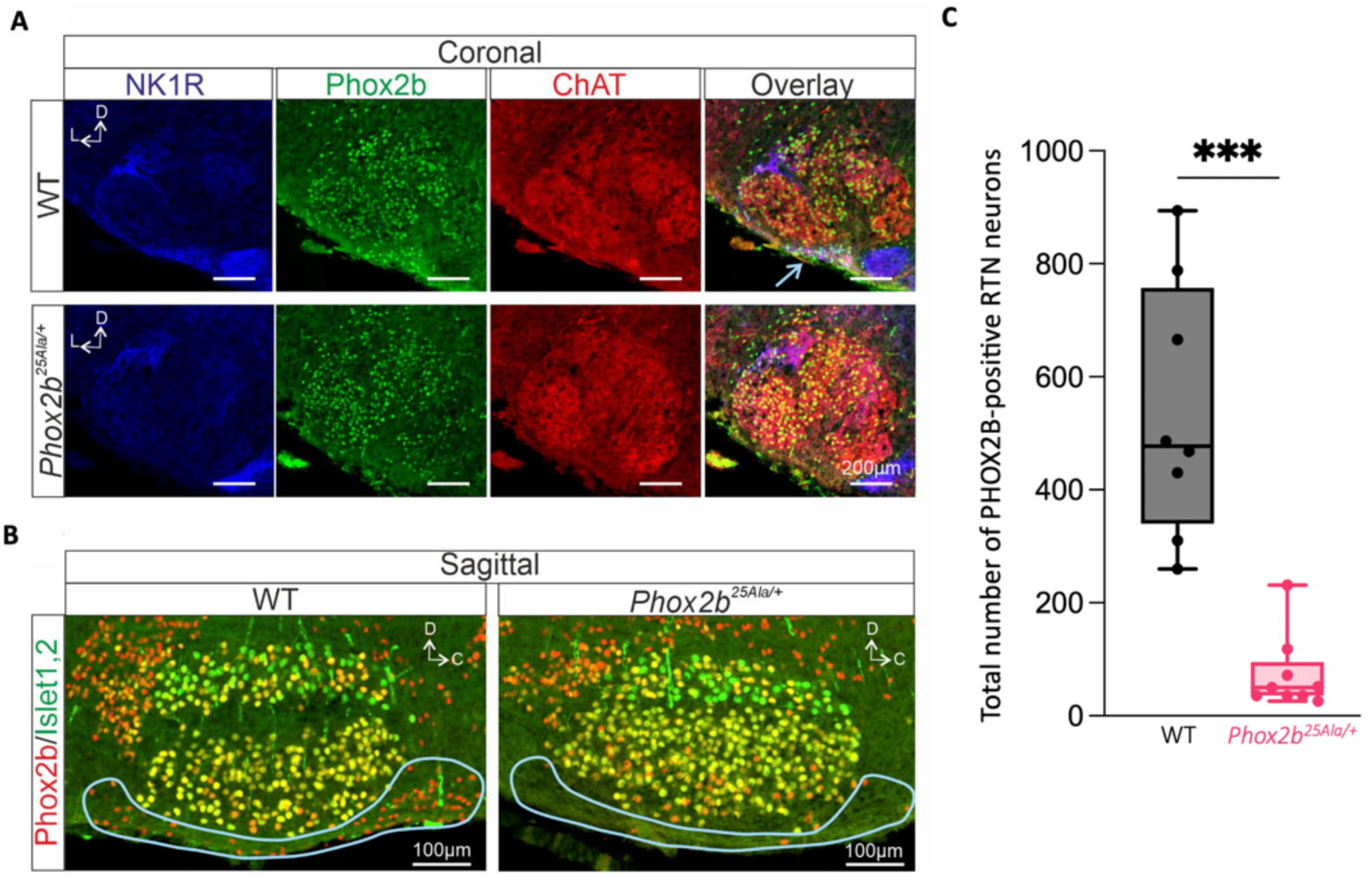
Dysgenesis of the RTN in constitutive *Phox2b^25Ala/+^* embryos. (A) Representative coronal sections through the facial motor nucleus (nVII) of E18.5 wild-type (WT; top) and constitutive *Phox2b^25Ala/+^* (bottom) embryos immunolabeled for NK1R (blue), PHOX2B (green) and ChAT (red). Merged images are shown in the right column. Arrows indicate PHOX2B-positive RTN neurons, which are abundant in WT embryos and markedly reduced in *Phox2b^25Ala/+^* embryos. (B) Representative sagittal sections through the nVII of E18.5 WT (left) and constitutive *Phox2b^25Ala/+^* (right) embryos stained for PHOX2B (red) and Islet1,2 (green). The RTN is delimited by the light blue line and contains markedly fewer PHOX2B-positive neurons in the mutant. (C) Quantification of PHOX2B-positive neurons detected in the RTN region for WT (n = 8, black) and *Phox2b^25Ala/+^* (n = 9, pink) embryos. Boxplots show the median, interquartile range, and full range, with individual animals represented by symbols. Two-sided exact Mann–Whitney test, *** *P* < 0.001.

### Surviving MafB-lineage mutants gain weight despite substantial neonatal mortality

To investigate the developmental origin of the phenotype observed in constitutive *Phox2b^25Ala/+^* mice, we generated *MafB::Phox2b^25Ala/+^*(*Phox2b^r5&6-25Ala/+^*) mice, in which the mutant allele is activated in MafB-lineage cells associated with the r5–r6 hindbrain territory. Postnatal survival and growth were assessed during the first week of life in *Phox2b^r5&6-25Ala/+^* mice (n = 15) and wild-type littermates (n = 14) from four litters, using the same monitoring protocol as for the constitutive model. Activation of the mutant allele in the MafB lineage did not restore neonatal survival to wild-type levels (*P* = 0.004; Fig. 7A). *Phox2b^r5&6-25Ala/+^* pups exhibited substantial mortality during the first two postnatal days, resembling that observed in constitutive *Phox2b^25Ala/+^* pups. At P2, 7 of 15 (46.7%) *Phox2b^r5&6-25Ala/+^* mice remained alive, compared with 13 of 14 (92.9%) wild-type littermates. Following one additional death between P2 and P3, the six surviving mutants remained alive through P7, whereas mortality continued beyond P3 in constitutive model.

**Figure 7.**
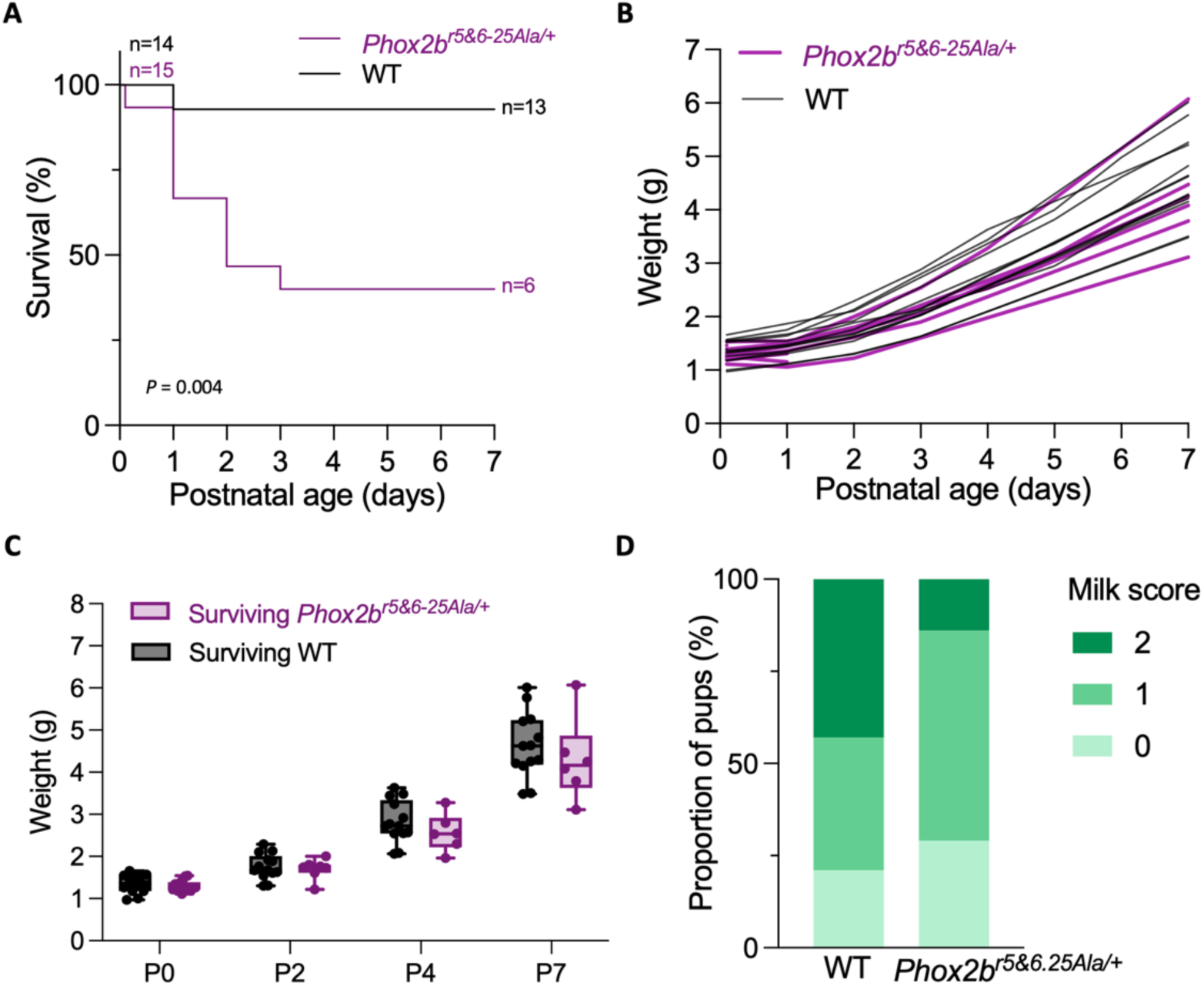
Neonatal lethality with preserved weight gain among surviving *MafB::Phox2b^25Ala/+^ (Phox2b^r5&6-25Ala/+^*) mice. (A) Kaplan–Meier survival curves of *Phox2b^r5&6-25Ala/+^* (n = 15, purple) mice and wild-type (WT) littermates (n = 14, black) from birth (P0) to postnatal day 7 (P7). Survival was significantly reduced in *Phox2b^r5&6-25Ala/+^* pups (two-sided log-rank test, *P* = 0.004). (B) Individual body weight trajectories of *Phox2b^r5&6-25Ala/+^* mice (n = 14, purple) and wild-type littermates (n = 14, black) from P0 to P7. Individual trajectories are discontinued after death. (C) Body weights of *Phox2b^r5&6-25Ala/+^* mice (purple) and wild-type littermates (black) alive at each indicated age. Body weight increased with age in both genotypes (age effect, *P* < 0.001), with no significant overall effect of genotype (*P* = 0.32) or genotype × age interaction (*P* = 0.73). Sample sizes for the mutant and WT groups, respectively, were n = 14 and 14 at P0, n = 7 and 13 at P2, and n = 6 and n = 13 at P4 and P7. Boxplots show the median, interquartile range, and full range, with individual animals represented by symbols. (D) Distribution of milk scores (0, no visible milk; 1, intermediate gastric milk content; and 2, milk-filled stomach) in wild-type (WT, n = 14) and *Phox2b^r5&6-25Ala/+^* (n = 14) pups at P0 (two-sided exact stratified Wilcoxon–Mann– Whitney test, stratified by litter, *P* = 0.051). Bars show the proportion of pups assigned to each score within each genotype.

Birth weight was comparable between wild-type and *Phox2b^r5&6-25Ala/+^*pups (1.35 ± 0.06 g vs. 1.30 ± 0.03 g, respectively; Holm-adjusted *P* = 0.50; Fig. 7C). Body weight increased throughout the first postnatal week in both genotypes, with no significant genotype × age interaction (*P* = 0.73; Fig. 7B,C). The estimated difference in weight-gain rate between mutant and wild-type pups was −0.058 g/day (95% CI, −0.147 to 0.031; *P* = 0.19). Analyses restricted to observed body weights yielded similar results. Thus, activation of the mutant allele in the MafB lineage was associated with substantial neonatal mortality, whereas postnatal weight gain was preserved among survivors, although these analyses do not establish equivalence between genotypes. This contrasting pattern suggests that the marked growth impairment observed in constitutive *Phox2b^25Ala/+^*mice involves PHOX2B-expressing populations that are not encompassed by, or are not affected to the same extent in, the MafB-lineage model.

### The reduction in gastric milk content is attenuated in MafB-lineage mutants compared with constitutive mutants

In the cohort used to assess postnatal survival and growth, milk-score distributions were numerically shifted toward lower values in *Phox2b^r5&6-25Ala/+^* pups than in wild-type littermates, although the difference did not reach statistical significance after accounting for litter (WT, n = 14; mutant, n = 14; *P* = 0.051; Fig. 7D). In the independent cohort studied by plethysmography, milk scores were significantly lower in *Phox2b^r5&6-25Ala/+^* pups than in wild-type littermates (WT, n = 32; mutant, n = 33; *P* = 0.013; Fig. 8).

**Figure 8.**
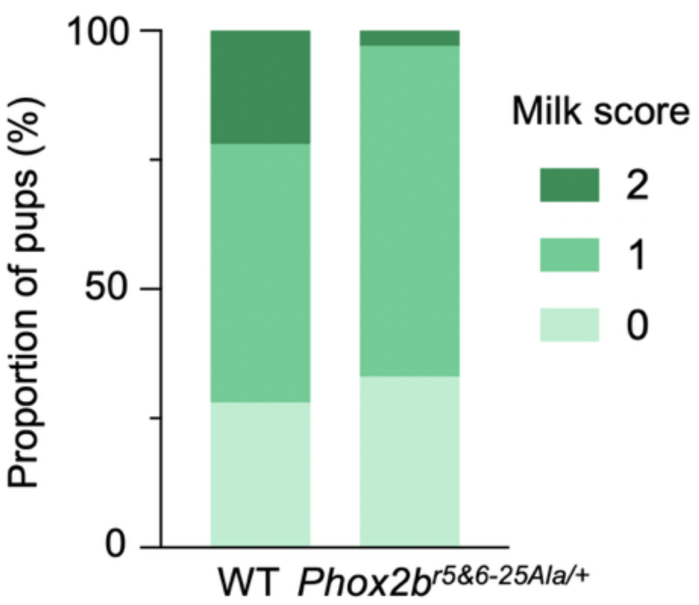
Gastric milk scores before plethysmography in wild-type (WT, n = 32) and *MafB::Phox2b^25Ala/+^* (*Phox2b^r5&6-25Ala/+^*, n = 33) pups at P0. Bars show the proportion of pups assigned to each milk score (0, no visible milk; 1, intermediate gastric milk content; and 2, milk-filled stomach) within each genotype. Milk-score distributions were shifted toward lower values in *Phox2b^r5&6-25Ala/+^* pups compared with WT littermates after accounting for litter (two-sided exact stratified Wilcoxon–Mann–Whitney test, stratified by litter, *P* = 0.013). All pups are shown; the statistical comparison included 27 WT and 33 mutant pups from litters containing both genotypes.

We next compared the genotype-associated shifts in milk-score distributions between the MafB-lineage and constitutive models across the four experimental cohorts. In a cumulative-link mixed model including experimental cohort as a fixed effect and litter as a random effect, milk scores were lower in constitutive *Phox2b^25Ala/+^* mutants than in their wild-type controls (odds ratio for a higher score, 0.03; 95% CI, 0.01–0.11; P < 0.001) and *Phox2b^r5&6-25Ala/+^* mutants than in their wild-type controls (odds ratio, 0.22; 95% CI, 0.08–0.62; P = 0.004). The reduction was significantly attenuated in the MafB-lineage model compared with the constitutive model (mutant × model interaction: β = 2.10; interaction odds ratio, 8.17; 95% CI, 1.70–39.13; P = 0.009). A linear mixed-model sensitivity analysis yielded a concordant result for the mutant × model interaction (P = 0.003). Thus, activation of the +5 allele in the MafB lineage attenuated, but did not abolish, the reduction in gastric milk scores associated with the mutant allele.

### MafB-lineage activation of the +5 mutation reproduces the major respiratory abnormalities of constitutive mutants

To examine the contribution of the MafB lineage associated with the r5**–**r6 hindbrain territory to the respiratory phenotype of constitutive *Phox2b^25Ala/+^* mice, we assessed ventilation in *Phox2b^r5&6-25Ala/+^*pups using the same protocol as for the constitutive model (nine litters; Fig. 9). In normoxia, *Phox2b^r5&6-25Ala/+^* mutant pups (n = 33) exhibited marked hypoventilation compared with wild-type littermates (n = 32; Fig. 9B) with ̇V_E_ reduced by 56% (*P* < 0.001). This reduction was driven by a 39% lower f_R_ (*P* < 0.001) and a 25% lower V_T_ (*P* < 0.001). Mutant pups exhibited a 2.8-fold higher apnea time than wild-type littermates (*P* < 0.001).

**Figure 9.**
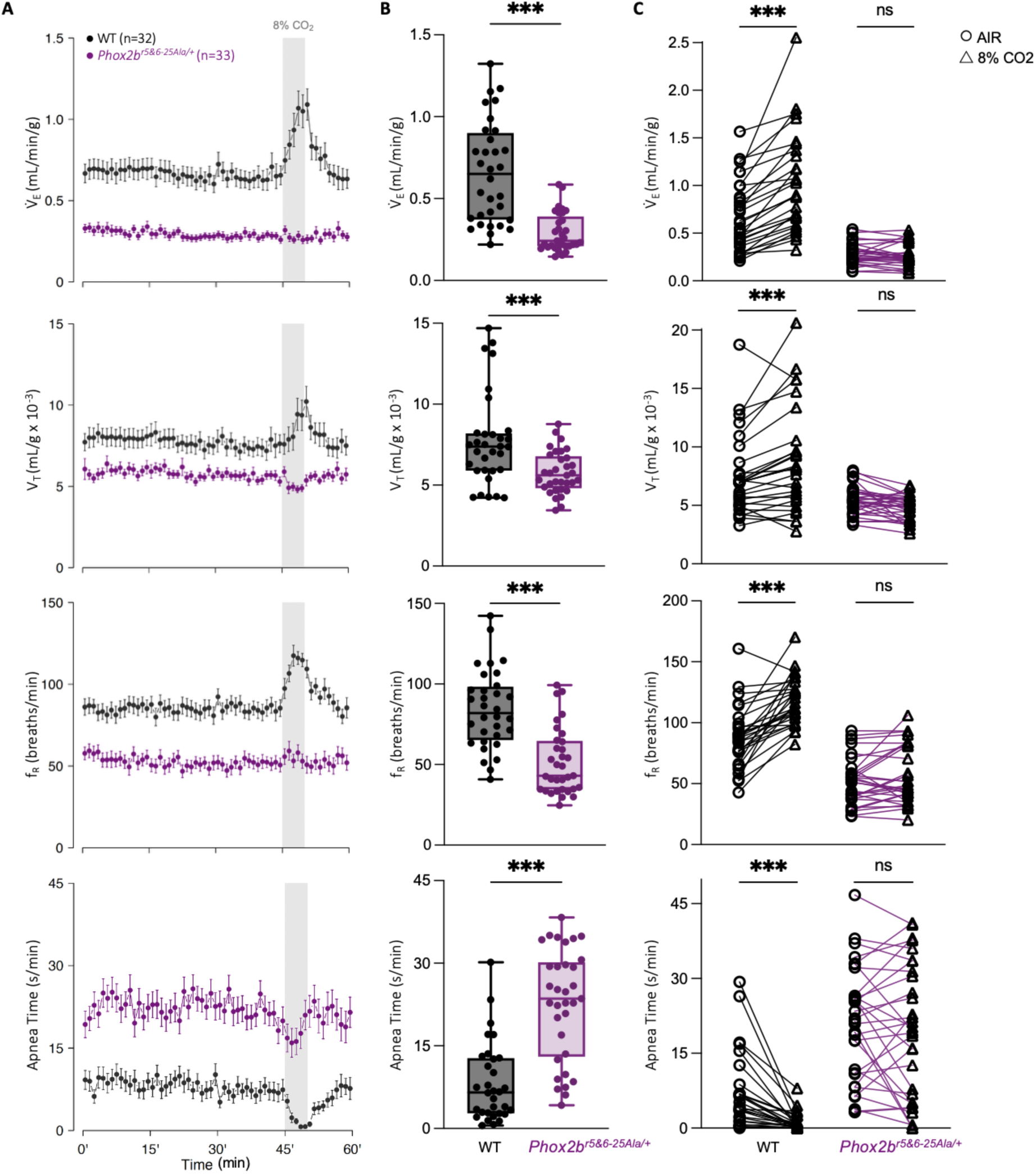
Depressed basal ventilation and blunted ventilatory response to hypercapnia in *MafB::Phox2b^25Ala/+^* (*Phox2b^r5&6-25Ala/+^*) mutants measured by whole-body plethysmography at 32°C. For each panel, minute ventilation (Vd_E_), tidal volume (V_T_), breathing frequency (f_R_), and apnea time are shown. (A) Ventilatory tracings (each dot represents mean ± SEM over a 1-min period) of *Phox2b^r5&6-25Ala/+^* mutant pups (n = 33, purple) and their wild-type (WT) littermates (n = 32, black) over the 60-min recording protocol, consisting of 45 min in room air, followed by 5 min of exposure to a gas mixture containing 8% CO₂ (gray shaded area), and 10 min of recovery in room air. (B) Baseline ventilation in *Phox2b^r5&6-25Ala/+^* mutants (n = 33, purple) compared with WT littermates (n = 32, black). Values were calculated during the first 40 min of exposure to room air. Boxplots show the median, interquartile range, and full range, with individual animals represented by symbols. (C) Individual ventilatory responses to hypercapnia in *Phox2b^r5&6-25Ala/+^* mutants (n = 31) and WT littermates (n = 28). Values were calculated during the 5 min of breathing room air preceding hypercapnic exposure (circles) and during the last 3 min of hypercapnic exposure (triangles). Paired measurements from individual pups are connected by lines (*Phox2b^r5&6-25Ala/+^*: purple; WT: black). Two-way mixed ANOVA revealed significant genotype x gas-condition interactions for ̇V_E_ (*P* < 0.001), V_T_ (*P* < 0.001), f_R_ (*P* < 0.001) and apnea time (*P* = 0.04). For all variables, Holm-Šídák-adjusted within-genotype comparisons showed significant changes during exposure to 8% CO₂ in WT pups (*P* < 0.001), whereas the corresponding changes were not significant in *Phox2b^r5&6-25Ala/+^* pups. Adjusted comparisons are indicated in the graphs. *** *P* < 0.001; ns, not significant.

Ventilatory responses to 8% CO₂ differed significantly between genotypes for all four ventilatory variables (genotype × gas-condition interactions: ̇V_E_, *P* < 0.001; f_R_, *P* < 0.001; V_T_, *P* < 0.001; apnea time, *P* = 0.04; Fig. 9C). In wild-type pups (n = 28), CO_2_ exposure increased ̇V_E_ by 52%, V_T_ by 18%, f_R_ by 34% and reduced apnea time by 83% (Holm-Šídák-adjusted *P* < 0.001 for all comparisons). By contrast, ̇V_E_ decreased slightly by 3% in *Phox2b^r5&6-25Ala/+^* pups (n = 31; adjusted *P* = 0.84), and no significant within-genotype changes were detected in f_R_, V_T_, or apnea time.

The RTN was also severely affected in *Phox2b^r5&6-25Ala/+^* embryos, with a near-complete loss of PHOX2B-positive neurons in the RTN region compared with wild-type littermates (two-sided unpaired Welch’s t test, *P* < 0.05; Fig. 10). This phenotype qualitatively resembled the severe RTN dysgenesis observed in constitutive *Phox2b^25Ala/+^* embryos. Together, these findings show that activation of the +5 allele in the MafB lineage produces baseline hypoventilation, increased apnea time, markedly impaired ventilatory responses to CO_2_ and severe dysgenesis of the RTN. Combined with the persistence of neonatal lethality, these results identify the MafB lineage as a major contributor to the respiratory and survival phenotypes of the constitutive +5 model.

**Figure 10.**
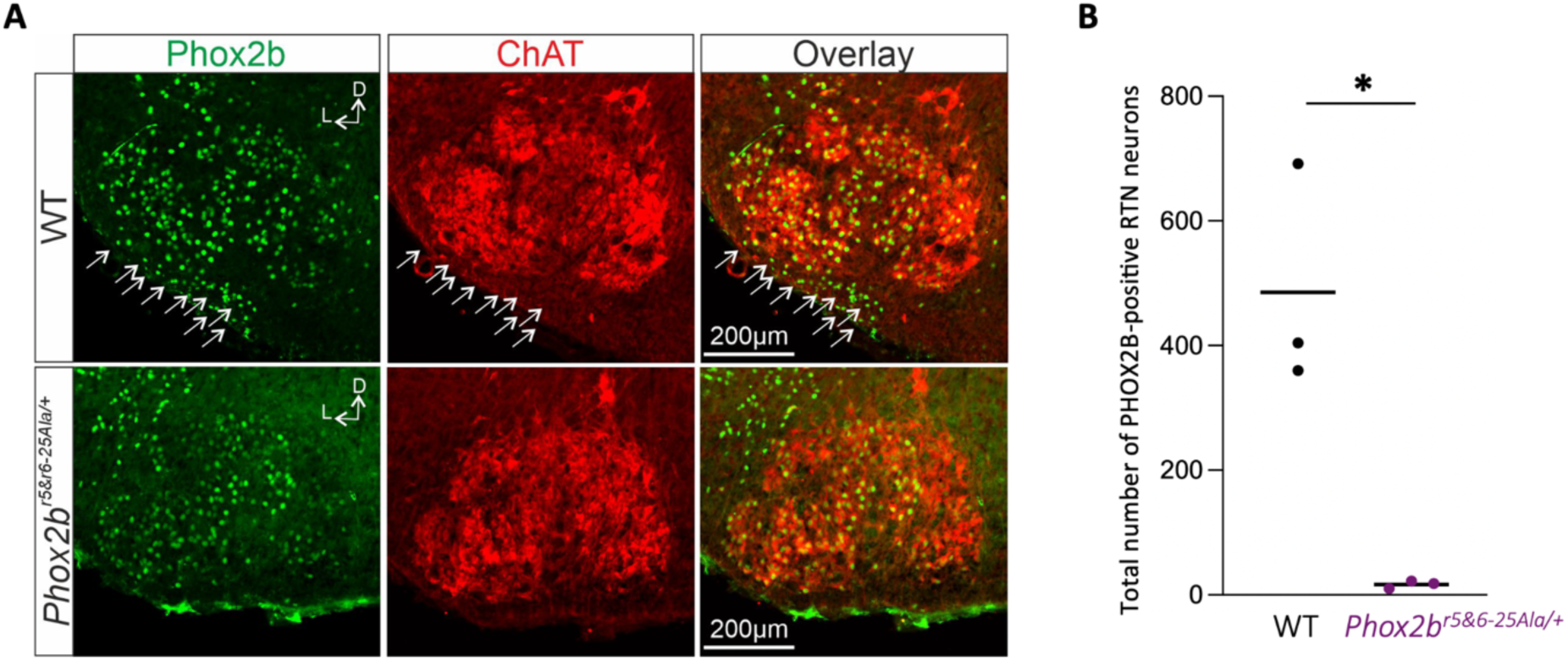
Dysgenesis of the RTN in *MafB::Phox2b^25Ala/+^* (*Phox2b^r5&6-25Ala/+^*) embryos. (A) Representative coronal sections through the facial motor nucleus (nVII) of E18.5 wild-type (WT; top) and conditional *Phox2b^r5&6-25Ala/+^* (bottom) embryos immunolabeled for PHOX2B (green) and ChAT (red). Merged images are shown in the right column. Arrows indicate PHOX2B-positive RTN neurons, which are abundant in WT embryos but markedly reduced in *Phox2b^r5&6- 25Ala/+^* embryos. (B) Total numbers of PHOX2B-positive neurons detected in the RTN region of WT (n = 3, black) and *Phox2b^r5&6-25Ala/+^* (n = 3, purple) embryos. Each point represents one embryo, and horizontal lines indicate group means. Groups were compared using a two-sided unpaired Welch’s t test (* *P* < 0.05).

## Discussion

In the present study, we generated a knock-in mouse model carrying the five-alanine expansion of the PHOX2B polyalanine tract associated with CCHS. Constitutive *Phox2b^25Ala/+^* pups exhibited major neonatal features of the disease, including hypoventilation, increased apnea time, markedly blunted ventilatory responses to hypercapnia, reduced respiratory-network activity and an impaired response to extracellular acidification *ex vivo*, and severe RTN dysgenesis. These abnormalities were accompanied by high neonatal mortality, reduced gastric milk scores, and markedly impaired postnatal weight gain among survivors. Activation of the mutant allele in the MafB lineage associated with the r5–r6 hindbrain territory also produced severe RTN dysgenesis, respiratory dysfunction, and early neonatal mortality. However, the reduction in milk scores was attenuated and postnatal weight gain was preserved among surviving MafB-lineage mutants. Together, these findings identify the MafB lineage as a major contributor to the respiratory and survival phenotypes caused by the +5 mutation and support partially separable developmental contributions to respiratory dysfunction and postnatal growth impairment.

Although polyalanine repeat mutations account for most cases of CCHS, their *in vivo* pathophysiology has previously been investigated mainly using the *Phox2b^27Ala/+^* model carrying a seven-alanine expansion^8–10^. This model established the developmental basis of impaired central chemosensitivity and revealed additional abnormalities affecting respiratory-network activity and upper-airway motor control. However, the death of constitutive +7 mutants within hours of birth limits the investigation of later postnatal manifestations. The +5 model described here was less rapidly lethal, with a subset of mutants surviving throughout the first postnatal week, while retaining the major respiratory features of CCHS. This phenotype is broadly consistent with the incomplete penetrance and clinical variability associated with the 20/25 genotype^1,6,7^. The *Phox2b^25Ala/+^* model therefore extends the available spectrum of PHOX2B PARM models and provides an opportunity to investigate early postnatal manifestations that cannot readily be studied in the rapidly lethal +7 model.

Despite its less rapidly lethal phenotype, the +5 mutation profoundly affected respiratory control from birth. Constitutive mutants exhibited marked baseline hypoventilation and a blunted ventilatory response to CO₂, with significant genotype × gas-condition interactions for minute ventilation, respiratory rate, and tidal volume, but not for apnea time. Respiratory-network dysfunction was already present at E18.5, when isolated mutant brainstem–spinal cord preparations exhibited a slower basal rhythm and an impaired response to extracellular acidification. These findings indicate that the postnatal respiratory phenotype arises, at least in part, from developmental abnormalities of central respiratory networks rather than solely from deterioration after birth. Although small residual increases in respiratory rate and minute ventilation were observed during CO₂ exposure, these changes were not statistically significant and were markedly smaller than those observed in wild-type pups. Their magnitude is compatible with the clinical variability in residual chemosensory responses, which is not strictly determined by PARM length^16,17^.

Severe respiratory abnormalities were also observed following activation of the +5 allele in the MafB lineage, including baseline hypoventilation, increased apnea time, and markedly impaired responses to CO₂. MafB-lineage mutants had only a small residual population of PHOX2B-positive RTN neurons, qualitatively resembling the marked RTN dysgenesis observed in constitutive mutants. This anatomical defect provides a plausible substrate for the impaired hypercapnic response and is consistent with lineage studies indicating that RTN neurons arise predominantly from r5-derived progenitors^12,18^. Early neonatal mortality also persisted, indicating that abnormalities affecting this lineage are sufficient to compromise survival. However, because the constitutive and MafB-lineage models were examined in separate respiratory and survival cohorts, differences in the magnitude of hypoventilation, CO₂ responsiveness, apnea time, or mortality between them should not be inferred from comparisons of significance levels or numerical estimates alone.

These findings expand upon those obtained with the previous conditional +7 model, in which Egr2/Krox20-Cre-mediated activation of the mutant allele targeted lineages derived from rhombomeres 3 and 5^10^. In that model, RTN development and the hypercapnic response were severely impaired, whereas neonatal survival was largely preserved, showing that loss of RTN-dependent chemosensitivity alone was insufficient to reproduce the lethality of constitutive +7 mutants. The present model targets the MafB lineage associated with r5 and r6 and combines RTN dysgenesis with hypoventilation, respiratory instability, and neonatal mortality. This pattern is consistent with evidence from the dB2 lineage that r5-derived populations contribute prominently to RTN development and the neonatal hypercapnic response, whereas r6-derived populations contribute to tidal-volume generation, respiratory stability, and survival^12^. Taken together, the parallels between these conditional models support, but do not demonstrate, a contribution of additional r6-derived PHOX2B populations to the respiratory and survival abnormalities observed in MafB-lineage mutants.

Postnatal growth showed a different pattern. Constitutive *Phox2b^25Ala/+^*survivors displayed markedly impaired weight gain, whereas surviving MafB-lineage mutants gained weight throughout the first postnatal week, with no detectable difference in weight trajectories compared with their wild-type littermates. This occurred despite substantial hypoventilation, impaired CO₂ responsiveness, increased apnea time, and early mortality in the MafB-lineage model. Gastric milk scores provided a complementary but more nuanced picture: scores were reduced in both mutant models, but this reduction was significantly attenuated in MafB-lineage mutants compared with constitutive mutants after adjustment for experimental cohort and litter. Thus, MafB-lineage activation attenuated, but did not abolish, the early reduction in gastric milk content, whereas the marked impairment of weight gain observed in constitutive mutants was not detected among surviving MafB-lineage mutants. Together, these findings suggest that severe respiratory dysfunction is not necessarily accompanied by marked postnatal growth impairment and that reduced gastric milk content alone does not fully account for the different growth patterns in the two models.

The mechanisms underlying these different growth patterns remain uncertain. Normal neonatal growth depends on effective milk acquisition and the coordination of suckling, swallowing, and breathing^19^. PHOX2B-expressing hindbrain populations contribute to premotor circuits involved in orofacial and ingestive behaviors, and populations not encompassed by the MafB-lineage model could contribute to the growth impairment observed in constitutive mutants^20,21^. However, gastric milk scoring provides only an indirect, single-time-point measure of feeding and cannot distinguish impaired suckling from altered swallowing, access to the dam, gastrointestinal dysfunction, or increased energy expenditure. The present data therefore do not identify the neuronal populations or physiological mechanisms responsible for the growth phenotype.

Several limitations should be considered. First, high neonatal mortality reduced the number of animals available for longitudinal measurements, and analyses restricted to survivors are susceptible to survival bias. Second, the constitutive and MafB-lineage models were studied in separate cohorts. Although milk scores could be compared using a common ordinal mixed model adjusted for experiment and litter, respiratory, growth, and survival outcomes were not analyzed together using common interaction models; quantitative equivalence between the two models should therefore not be inferred. Third, MafB-Cre targets a broad developmental lineage that is not confined to the hindbrain. Although its hindbrain activity encompasses cells associated with the r5–r6 territory, the present experiments do not distinguish the contributions of r5, r6, individual neuronal populations or extra-hindbrain MafB-lineage cells. Fourth, gastric milk scoring is an indirect and relatively coarse measure of neonatal feeding, and the mechanisms responsible for impaired postnatal growth were not directly examined. Finally, genetic background, litter effects, and other modifiers may influence the penetrance and severity of PHOX2B polyalanine-expansion phenotypes in both mice and patients.

In conclusion, the *Phox2b^25Ala/+^* mouse reproduces major neonatal respiratory features of CCHS while allowing a subset of mutants to survive beyond the period accessible in the established constitutive +7 model. Activation of the mutant allele in the MafB lineage is sufficient to produce severe RTN dysgenesis, respiratory dysfunction, and neonatal mortality, whereas postnatal weight gain is preserved among survivors. These findings support partially separable developmental contributions to the respiratory and growth phenotypes produced by the +5 mutation and provide a model for investigating early postnatal disease mechanisms and for the future evaluation of candidate interventions.

## Acknowledgements

We thank Dr. L. Hernandez-Miranda (Institut für Molekulare und Zelluläre Anatomie, Universität Ulm) for providing *MafB*Cre mouse line, Dr. J.-F. Brunet (Institut de Biologie de l’École Normale Supérieure) for his advice on the genetic strategy, Dr. M.-C. Birling (Institut de Génétique et de Biologie Moléculaire et Cellulaire, CNRS and INSERM) and the PHENOMIN-iCS team for the generation of the conditional *Phox2b^25Ala^*mouse line, Dr. G. Morera (Centre Régional d’Exploration Fonctionnelle et Ressources Expérimentales, Université Paul Sabatier and INSERM) for rederiving the *Phox2b^25Ala^* mouse line by *in vitro* fertilization using cryopreserved sperm, and Dr. A. Diet (TAAM-PHENOMIN-UAR44, CNRS) and the TAAM team for rederiving the *MafB*Cre mouse line from cryopreserved embryos.

## Author contributions

N.Ra., C.R. and A.M. performed most of the research and equally contributed to this work. M.T.-B. and B.M. supervised the study and equally contributed to this work. Conceptualization: J.G., B.D., P.B., S.D., M.-P.O., C.D., M.T.-B. and B.M. Investigation: N.Ra., C.R., A.M., P.A., T.B., J.-L.M., L.C., J.F., N.Ro., M.P., N.A., E.S., and M.R. Methodology: N.Ra., C.R., A.M., T.B., M.R., M.T.-B. and B.M. Visualization: N.Ra., C.R., A.M., P.A., J.-L.M., L.C., J.F., N.Ro., M.P., N.A., M.T.-B. and B.M. Validation: N.Ra., C.R., A.M., P.A., T.B., J.-L.M., L.C., J.F., M.R., M.T.-B. and B.M. Formal analysis: N.Ra., C.R., A.M., P.A., J.G., B.D., P.B., S.D., M.-P.O., C.D., M.T.-B. and B.M. Data curation: N.Ra., C.R., A.M., P.A., J.-L.M., L.C., J.F., N.Ro., M.T.-B. and B.M. Software: N.Ra., C.R., A.M., M.T.-B. and B.M. Resources: A.M., P.A., T.B., L.C., M.R., M.T.-B. and B.M. Funding acquisition: S.D., M-P.O., C.D., M.T.-B. and B.M. Writing—original draft: M.T.-B. and B.M., with input from all coauthors. Writing— review and editing: M.T.-B. and B.M., with input from all coauthors.

## References

1. Trang, H. et al. Guidelines for diagnosis and management of congenital central hypoventilation syndrome. Orphanet J Rare Dis 15, 252 (2020).

2. Dudoignon, B. et al. European central hypoventilation syndrome consortium description of congenital central hypoventilation syndrome neonatal onset. Eur J Pediatr 184, 161 (2025).

3. Pattyn, A., Goridis, C. & Brunet, J. F. Specification of the central noradrenergic phenotype by the homeobox gene Phox2b. Mol Cell Neurosci 15, 235–43 (2000).

4. Pattyn, A., Morin, X., Cremer, H., Goridis, C. & Brunet, J. F. The homeobox gene Phox2b is essential for the development of autonomic neural crest derivatives. Nature 399, 366–70 (1999).

5. Dauger, S. et al. Phox2b controls the development of peripheral chemoreceptors and afferent visceral pathways. Development 130, 6635–42 (2003).

6. Weese-Mayer, D. E., Rand, C. M., Zhou, A., Carroll, M. S. & Hunt, C. E. Congenital central hypoventilation syndrome: a bedside-to-bench success story for advancing early diagnosis and treatment and improved survival and quality of life. Pediatr Res 81, 192– 201 (2017).

7. Bachetti, T. & Ceccherini, I. Causative and common PHOX2B variants define a broad phenotypic spectrum. Clin Genet 97, 103–113 (2020).

8. Dubreuil, V. et al. A human mutation in Phox2b causes lack of CO2 chemosensitivity, fatal central apnea, and specific loss of parafacial neurons. Proc Natl Acad Sci U S A 105, 1067–72 (2008).

9. Madani, A. et al. Obstructive Apneas in a Mouse Model of Congenital Central Hypoventilation Syndrome. Am J Respir Crit Care Med 204, 1200–1210 (2021).

10. Ramanantsoa, N. et al. Breathing without CO(2) chemosensitivity in conditional Phox2b mutants. J Neurosci 31, 12880–8 (2011).

11. Hernandez-Miranda, L. R. et al. Mutation in LBX1/Lbx1 precludes transcription factor cooperativity and causes congenital hypoventilation in humans and mice. Proc Natl Acad Sci U S A 115, 13021–13026 (2018).

12. Cui, K. et al. Genetic identification of medullary neurons underlying congenital hypoventilation. Sci. Adv. 10, eadj0720 (2024).

13. Birling, M.-C., Dierich, A., Jacquot, S., Hérault, Y. & Pavlovic, G. Highly-efficient, fluorescent, locus directed cre and FlpO deleter mice on a pure C57BL/6N genetic background. Genesis 50, 482–489 (2012).

14. Wu, X. et al. Mafb lineage tracing to distinguish macrophages from other immune lineages reveals dual identity of Langerhans cells. J Exp Med 213, 2553–2565 (2016).

15. Matrot, B. et al. Automatic classification of activity and apneas using whole body plethysmography in newborn mice. J Appl Physiol 98, 365–70 (2005).

16. Carroll, M. S. et al. Residual chemosensitivity to ventilatory challenges in genotyped congenital central hypoventilation syndrome. J Appl Physiol (1985) 116, 439–50 (2014).

17. Bokov, P. et al. Central CO_2_ chemosensitivity and CO_2_ controller gain independently contribute to daytime P CO_2_ in young subjects with congenital central hypoventilation syndrome. Journal of Applied Physiology 135, 343–351 (2023).

18. Dubreuil, V. et al. Defective respiratory rhythmogenesis and loss of central chemosensitivity in Phox2b mutants targeting retrotrapezoid nucleus neurons. J Neurosci 29, 14836–46 (2009).

19. Maynard, T. M., Zohn, I. E., Moody, S. A. & LaMantia, A.-S. Suckling, Feeding, and Swallowing: Behaviors, Circuits, and Targets for Neurodevelopmental Pathology. Annu Rev Neurosci 43, 315–336 (2020).

20. Dempsey, B. et al. A medullary centre for lapping in mice. Nat Commun 12, 6307 (2021).

21. Sungeelee, S., et al. A genetically defined pontine nucleus essential for ingestion in mice. Proc. Natl. Acad. Sci. U.S.A. 122, e2411174122 (2025).

